# Immediate glucose signaling transmitted via the vagus nerve in gut-brain neural communication

**DOI:** 10.1101/2024.03.27.586971

**Authors:** Serika Yamada, Akiyo Natsubori, Kazuki Harada, Takashi Tsuboi, Hiromu Monai

## Abstract

Sucrose consumption is influenced by certain gut-brain signaling mechanisms. Among these, one pathway involves neuropod cells, which form synaptic connections with the vagus nerve, leading to the immediate activation of central dopaminergic pathways. This study explored the role of the frontal cortex in its process. We found that the vagus nerve’s immediate activation is mediated by the sodium-glucose cotransporter 1 (SGLT1) of neuropod cells after the intragastric glucose injection in mice. Also, we showed that the involvement of both astrocytes and neurons in the frontal cortex via D2 and D1 dopamine receptors, respectively, by in vivo Ca^2+^ imaging. Finally, we revealed that psychological stress, which induces a reduction in sucrose preference, significantly diminishes the activation levels of both the vagus nerve and the frontal cortex. These findings highlight the role of a comprehensive gut-brain network in modulating sucrose preference, involving neuropod cells, the vagus nerve, and the frontal cortex.

**Graphical abstract:** 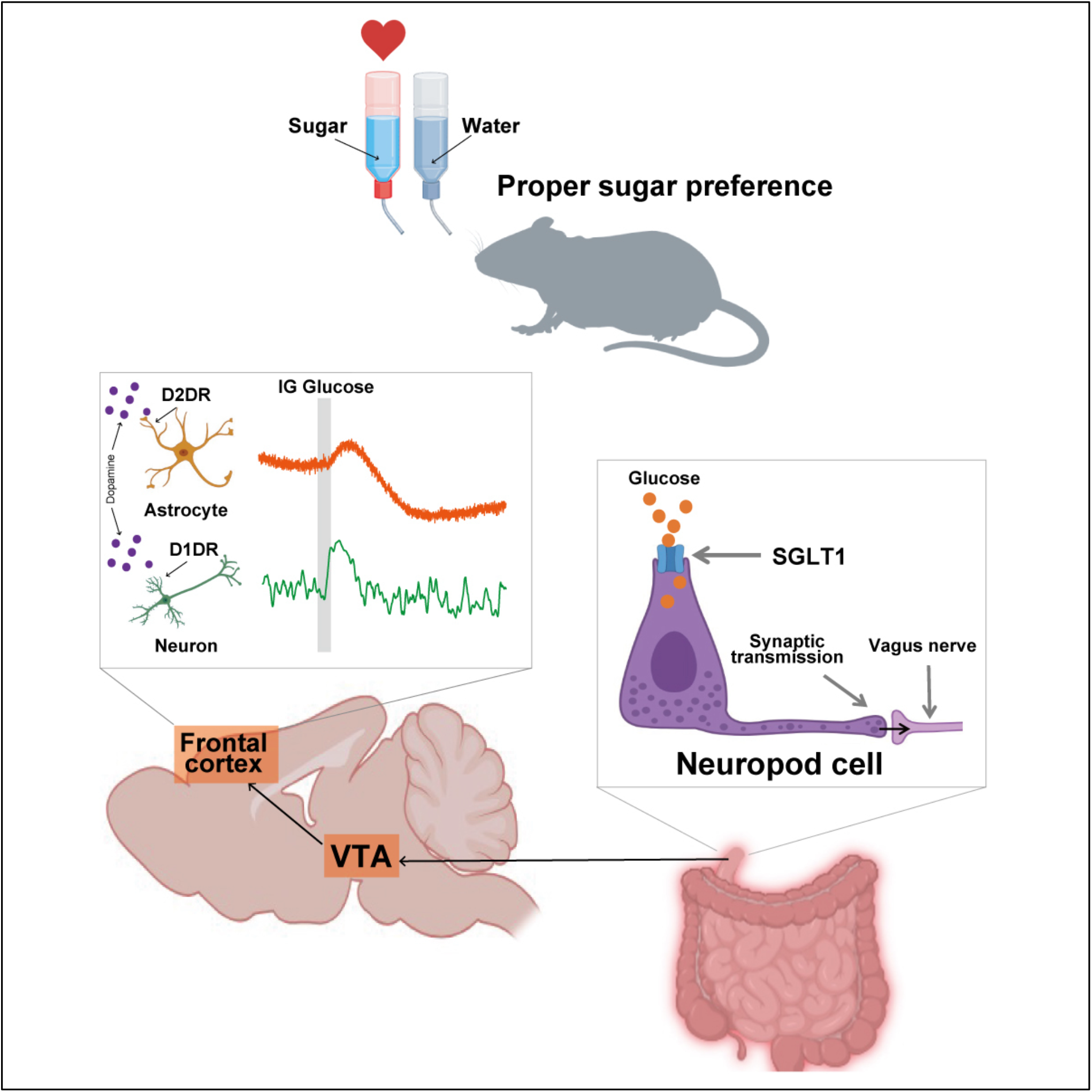

## Introduction

Glucose is one of the primary source of energy for animals. Though it is ingested orally, in recent years, the importance of signals via the vagus nerve, which afferently projects from the intestine to the brain, following glucose intake in regulating sugar (glucose or sucrose) consumption has garnered considerable academic interest.^1–6^ Central regulation of sugar intake via the afferent vagus nerve is conducted in complex pathways involving both inhibitory and stimulatory mechanisms, attributed to its difference of projection target corresponding to vagus nerve cell types.^7–10^ The representative example of the feeding inhibitory pathway mediated by the vagus nerve is gastrointestinal hormones (ex. GLP-1), which ultimately regulate feeding volume by modulating like NPY/AgRP neurons and POMC/MSH in the arcuate nucleus and paraventricular nucleus.^7,11–17^ While the feeding inhibitory mechanism operates via the vagus nerve and is mediated by the hypothalamus, the promotion of feeding through the vagus nerve involve interaction with dopaminergic neurons in other brain regions to which they project. Both pathways are crucial for maintaining sucrose preference.^18–21^ Recent findings have revealed that activation of the vagus nerve following glucose intake stimulates both pathways. Additionally, it has been suggested that activation of the right vagus nerve primarily activates the Nigrostriatal dopaminergic pathway, whereas activation of the left vagus nerve predominantly activates the Ventral Tegmental Area (VTA), which is the originating region of the Mesocortical and Mesolimbic dopaminergic pathway.^2,3,7,22–27^ By these complex regulations, sucrose preference of healthy mice in a two-bottle choice test consistently exceeds 80%. Also, it is known that mice subjected to chronic psychological stress exhibit a marked decline in sucrose preference because of decrease in motivational drive for sucrose consumption and a reduced perception of pleasure post-sucrose intake.^28–31^

Gastrointestinal hormones contribute not only to feeding inhibition but also to the promotion of feeding. Various gastrointestinal hormones are secreted minutes after glucose uptake in the intestine, with each hormone activating either the left or right vagus nerve. Consequently, several minutes after glucose reaches the intestine, activation of the nigrostriatal pathway via the right vagus nerve and activation of the mesolimbic pathway via the left vagus nerve leads to dopamine release in the dorsal and ventral striatum, respectively, which promotes feeding.^7,23,32^ As mentioned above, the dopamine release in the dorsal and ventral striatum induced by such digestive hormones occurs minutes after glucose uptake in the intestine. However, recent observations have shown that dopaminergic neurons in the VTA, originating from the left vagus nerve, are activated within seconds of administering sucrose into the intestine.^22^

In recent years, neuropod cells, a subtype of enteroendocrine cells, have been identified as immediate activators of the vagus nerve under intragastric (IG) sugar injection. These cells are predominantly situated in the duodenum, where they establish synaptic connections with the vagus nerve. Neuropod cells assimilate glucose through sodium-glucose co-transporter 1 (SGLT1) and subsequently activate the vagus nerve within a mere second by discharging glutamate as a neurotransmitter. Signals derived from neuropod cells have been demonstrated to contribute to the sucrose preference in mice.^33–38^

The frontal cortex is a principal projection target of VTA dopaminergic neurons. This dopaminergic mesocortical pathway is activated by reward responses and plays a role in prompting appropriate behavior in response to rewarding stimuli.^18, 39–45^ While the projection targets of VTA dopaminergic neurons encompass both the ventral striatum and the frontal cortex, a significant temporal divergence is observed: dopamine release in the ventral striatum occurs several minutes post-intestinal glucose injection, whereas VTA dopaminergic neurons are activated promptly within seconds. This disparity suggests that the immediate activation of VTA dopaminergic neurons under intestinal glucose injection might involve the frontal cortex through a pathway distinct from that associated with the striatum. Yet, it remains to be elucidated whether signals originating from neuropod cells activate neurons or astrocytes in the frontal cortex immediately.

While chronic psychological stress is implicated in dysregulating the brain’s reward system, recent reports have also underscored its influence on vagus nerve functionality.^46–53^ Given the role of vagus nerve mediated dopaminergic neuron activation in motivation and pleasure, it is postulated that signals from neuropod cells may play a role in the attenuated sucrose preference observed post-stress, although this hypothesis remains to be rigorously examined.

Building on existing literature, this study examines the immediate activation of frontal cortex by neuropod cell-mediated responses under direct IG glucose injection by performing transcranial cortex-wide Ca^2+^ imaging and electrophysiological vagus nerve recordings, then we investigated the signal complication by performing fiber photometric recording and two-photon Ca^2+^ imaging. Furthermore, we probed the potential modification in the neuropod cell-derived signaling in mice subjected to chronic mild restraint stress (CMRS).

## Results

### Immediate activation of the vagus nerve under intragastric (IG) glucose injection is mediated by neuropod cells

To investigate the vagus nerve’s rapid response via neuropod cells to direct glucose delivery into the duodenum, a 10% glucose solution was infused into the proximal duodenum of mice under anesthesia with 0.8-1.0% isoflurane. A pre-attached catheter facilitated the glucose solution’s injection over 8 seconds (**Fig. 1A**, see method). This procedure resulted in a notable activation of the left vagus nerve within 1 second of IG glucose injection (**Figs. 1B**, top and **C**, top), in stark contrast to the absence of such activation under IG water injection (**Figs. 1B**, bottom, **C**, bottom, and **D**, water vs. glucose: 0.96 ± 0.14 vs. 2.81 ± 1.56, p = 0.0043).

**Fig. 1.**
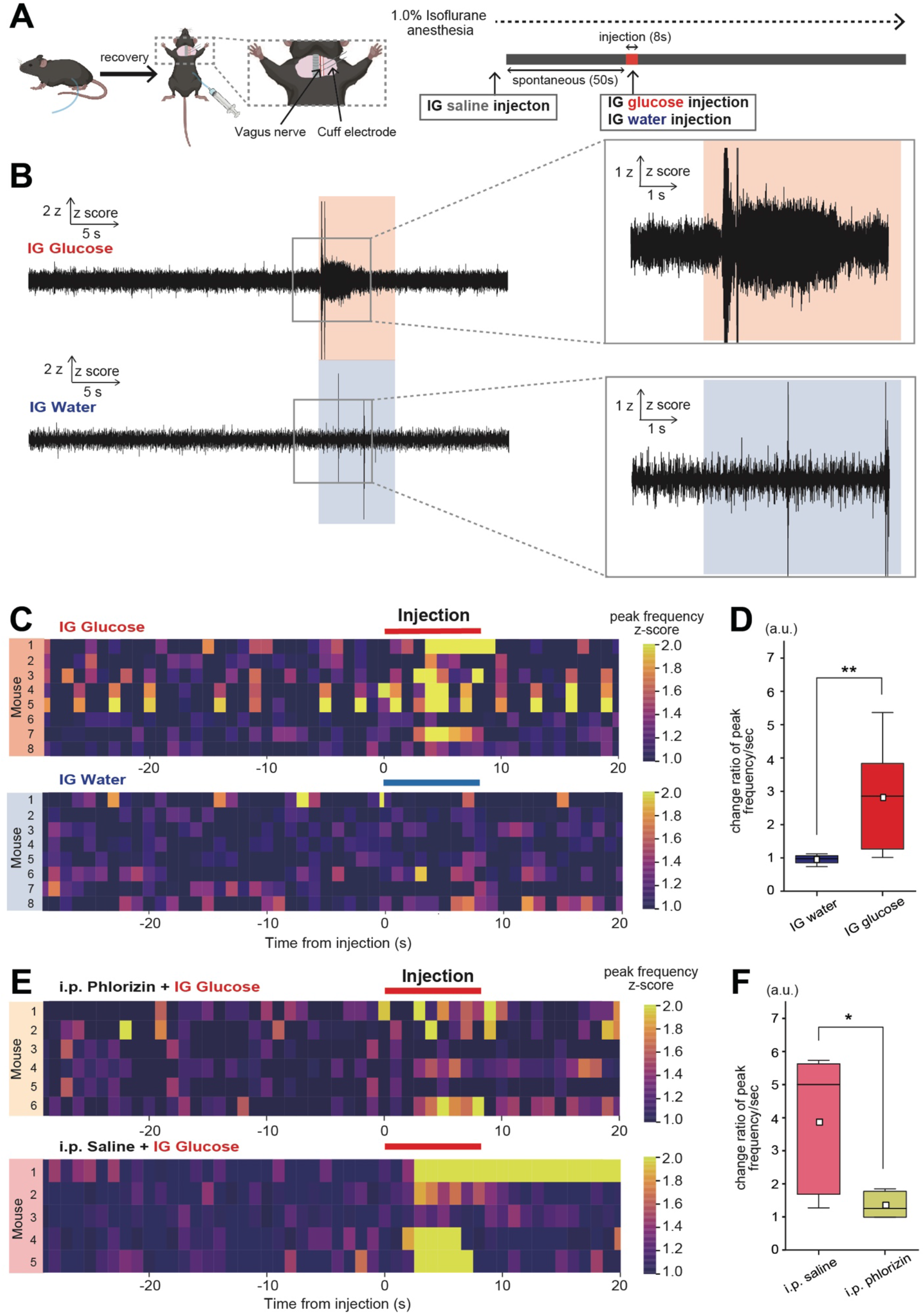
Vagal nerve response dynamics to IG glucose and water injections. (A) Sequential representation of vagus nerve recordings procedures during IG injection. (B) Illustration of 50Hz high-pass filtered vagus nerve activity under IG glucose (upper) and water (lower) injections. The periods of injection are highlighted in light red and light blue, respectively. (C) Graph depicting the normalized frequency of peaks per second in the vagus nerve activity of individual subjects from the IG glucose (upper) and water (lower) injection cohorts. Normalization was based on the average baseline in the spontaneous state (30 seconds prior to IG injection). (D) Comparative analysis of changes in the frequency of peaks per second in the vagus nerve activity in response to the IG water injection cohort (blue) and the IG glucose injection cohort (red). The initial 30 seconds prior to IG injection are designated as the pre-stimulation phase, and the 2-4 seconds after IG injection as the post-stimulation phase. **p<0.01 (IG glucose; N=9 mice, IG water; N=8 mice, two-sample t-test) (E) Graph illustrating the normalized frequency of peaks per second in the vagus nerves activity of individual subjects from the IG glucose cohort following i.p. phlorizin injection (upper data) and IG glucose cohort following i.p. saline injection (lower data). Normalization was performed based on the average baseline in the spontaneous state (30 seconds prior to IG injection). (F) Comparative analysis of changes in the frequency of peaks per second in the vagus nerve activity in response to the IG glucose injection cohort post-i.p. phlorizin injection (pink) and the IG glucose injection cohort post-i.p. saline injection (yellow). The initial 30 seconds prior to IG injection are designated as the pre-stimulation phase, and the 2-4 seconds after IG injection as the post-stimulation phase. **p<0.05 (IG glucose injection post-i.p. saline injection; N=5 mice, IG glucose injection post-i.p. phlorizin injection; N=6 mice, two-sample t-test)

The swift propagation of signaling is postulated to involve neuropod cells, a specialized type of intestinal endocrine cell. Neuropod cells integral to glucose assimilation via sodium-glucose co-transporter 1 (SGLT1)^33–38^. Subsequent experiments employing the SGLT1 inhibitor phlorizin demonstrated that mice did not exhibit significant vagus nerve activation (**Fig. 1E**, top, bottom and **1F**, saline vs. phlorizin: 2.20 ± 0.98 vs. 0.39 ± 0.16, p = 0.02).

Given that neuropod cells are presently the sole identified agents inducing immediate (within 1 second of glucose absorption) vagus nerve activation that leads sugar preference post-intestinal glucose absorption,^22, 34, 36^ the immediate response within 1 second of IG glucose injection implicates a mechanism originating from neuropod cells.

### The frontal cortex undergoes immediate activation under IG glucose injection

To assess the effect of IG glucose injection on cortical activity, transcranial cortex-wide Ca^2+^ imaging was conducted (**Fig. 2A**). This technique employed G7NG817 line transgenic mice, characterized by the expression of the Ca^2+^ sensor protein G-CaMP7, under the control of the GLT-1 promoter that express in astrocytes and excitatory neurons.^54,55^ Under the isoflurane anesthesia, the cortical Ca^2+^ spontaneous activity occurred regularly. This Ca^2+^ activity was synchronized across intracortical regions, with relatively high amplitudes in the medial occipital regions.^56–58^ In contrast, the frontal cortex exhibited marked activation immediately after the start of the IG glucose injection (**Fig. 2A**, middle), but not after the start of the IG water injection (**Figs. 2A**, bottom, and **B**).

**Fig. 2.**
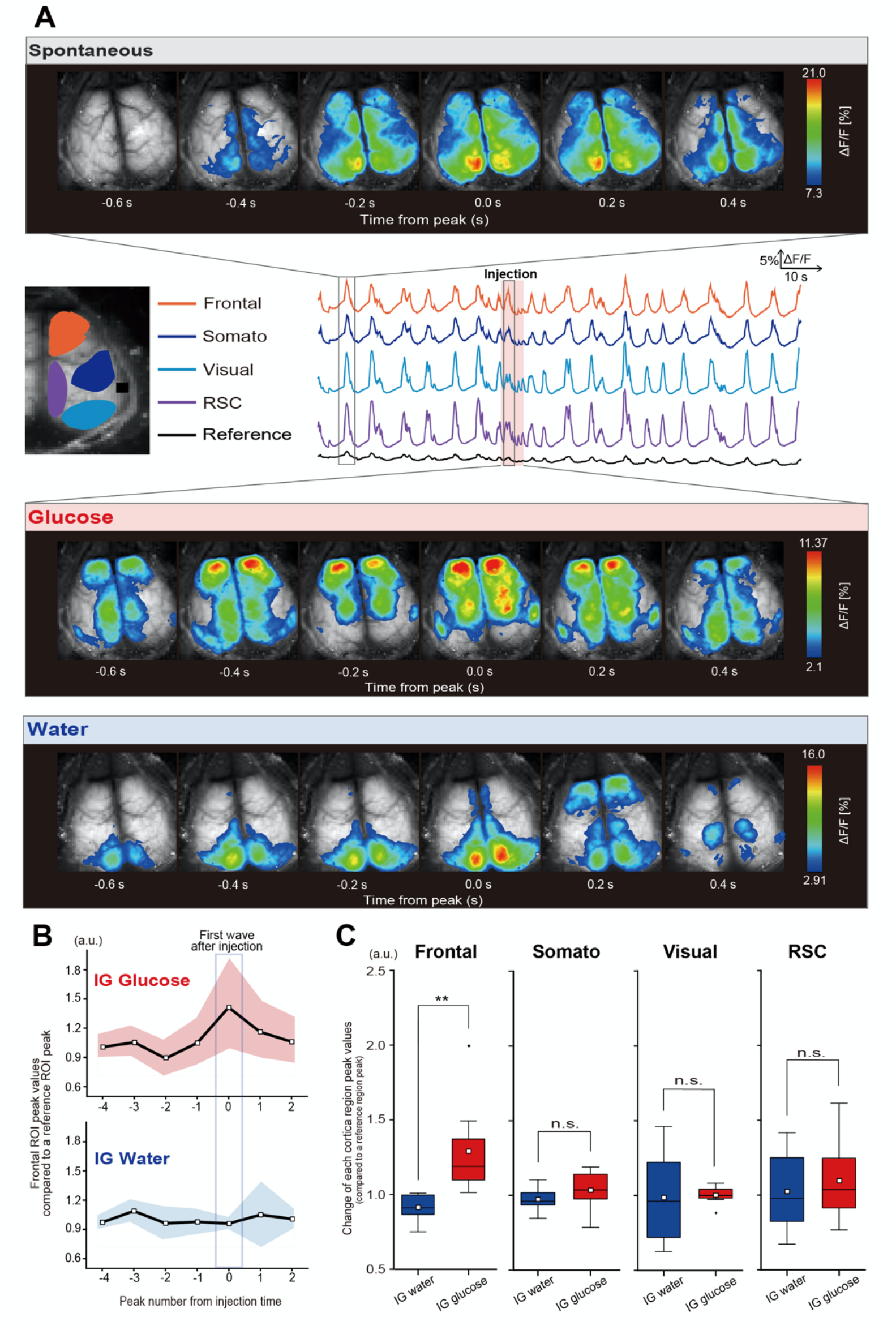
Differential cortical Ca^2+^ responses to IG glucose and water injection. (A) Illustration of representative fluorescence intensity (ΔF/F) traces in G7NG817 mice for each cortical region (Frontal: orange, Somato: blue, Visual: light blue, RSC: purple, Reference: gray). Upper panel depicts the cortical spontaneous Ca^2+^activity patterns. Middle and lower panels depict the cortical activity patterns under IG glucose and IG water injection, respectively. The analysis targets the earliest Ca^2+^ wave appearing within 3–8 seconds after the injection start. The pseudocolor representation employs the peak of the Ca^2+^ transient as the maximum value - 1SD, and the mean + 1SD as the minimum value. (B) Temporal variation of peak value of each Ca^2+^ wave in the Frontal ROI under the IG glucose (upper) and IG water (lower) injection. Data are normalized by the corresponding Ca^2+^ peak value in the reference ROI. The colored areas indicate the average ± SEM. (C) Comparison of Ca^2+^ activation levels under IG glucose and water injection in each cortical region. The data shown the peak value that appeared earliest between 3-8 seconds after the start of the injection relative to the average of the peak value during the 50 seconds prior to the injection. (IG glucose; N = 8 mice, IG water; N = 8 mice, two-sample t-test)

Quantitative analysis revealed that the IG glucose injection-induced Ca^2+^ activation was significant in the frontal region (**Fig. 2C**, Frontal: Water vs. Glucose: 0.92 ± 0.08 vs. 1.30 ± 0.30, p = 0.0063, Somato: Water vs. Glucose: 0.97 ± 0.072 vs. 1.04 ± 0.12, p = 0.27, Visual: Water vs. Glucose: 0.99 ± 0.28 vs. 1.00 ± 0.06, p = 0.89, and RSC: Water vs. Glucose: 1.02 ± 0.25 vs. 1.10 ± 0.26, p = 0.60). These results suggest that the unique responsiveness of the frontal cortex to IG glucose injection, signifying its immediate activation.

### Neuropod cells facilitate the immediate activation of the frontal cortex under IG glucose injection

In Figure 1, we observed a significant reduction in the immediate activation of the left vagus nerve under IG glucose injection in the presence of phlorizin, an inhibitor against SGLT1 crucial for glucose uptake by neuropod cells. Base on the results, to reveal the neuropod cell’s contribution to frontal cortex activation of IG glucose injection, we performed similar transcranial cortex-wide Ca^2+^ imaging in the presence of phlorizin (**Fig. 3A**).

**Fig. 3.**
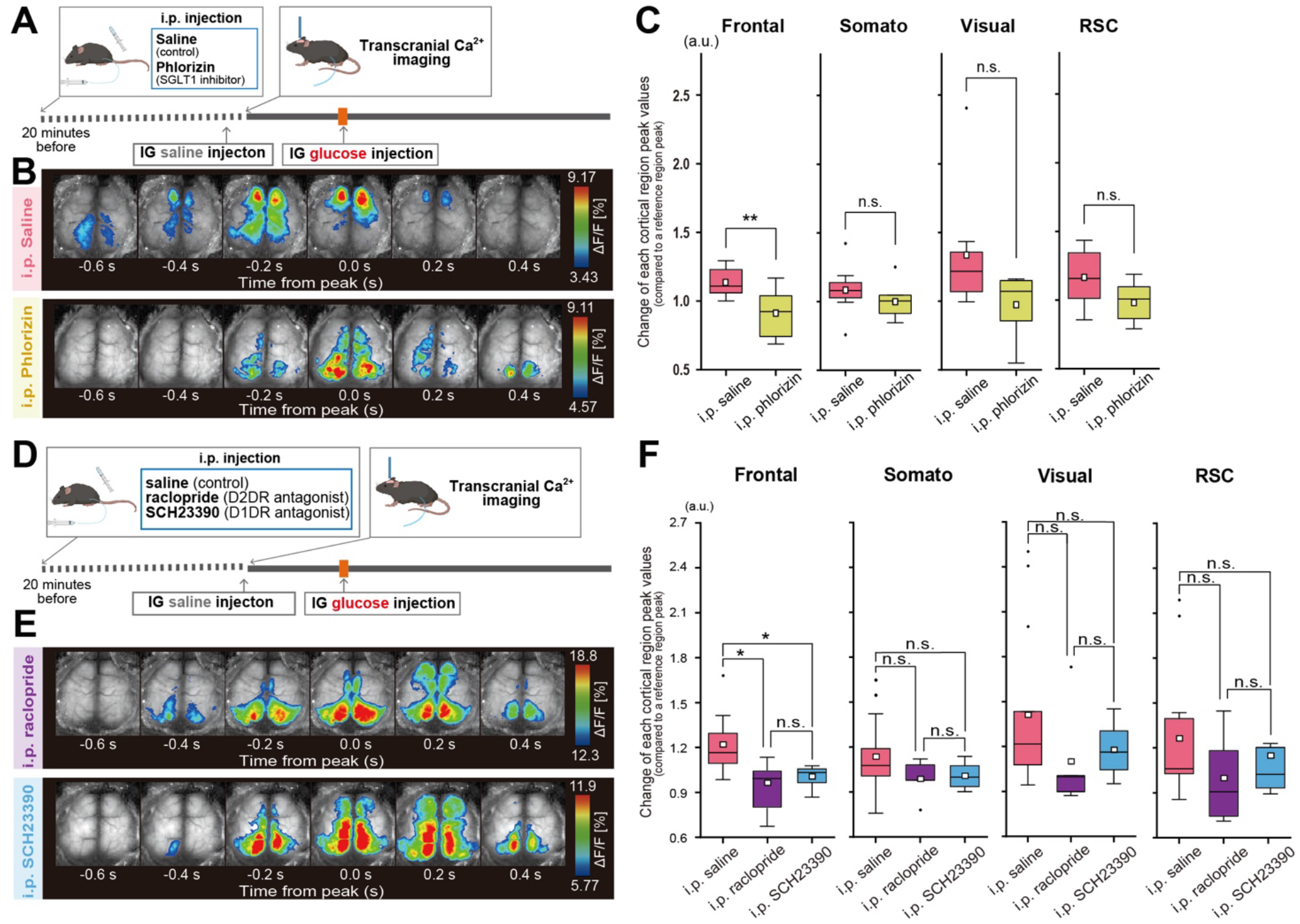
Influence of SGLT1 and dopaminergic antagonists on cortical Ca^2+^ activation under IG glucose injection. (A) Sequence of transcranial cortex-wide Ca^2+^ imaging under IG glucose injection with phlorizin (SGLT1 antagonist) and saline (control) administered intraperitoneally. (B) Illustration of cortical activity patterns under IG glucose injection with saline (upper) and with phlorizin (lower). The analysis targeted the earliest Ca^2+^ wave appearing within 3–8 seconds after injection. The pseudocolor representation employs the peak of the Ca^2+^ transient as the maximum value - 1SD and the mean + 1SD as the minimum value. (C) Comparison of Ca^2+^activation levels in each cortical region under IG glucose injection with saline and phlorizin pretreatment. The methodology entailed comparing the peak fluorescence intensity values of each brain region against those of a reference region (auditory cortex), encompassing the 50 seconds prior to injection and the subsequent post-injection period. Post-injection values were normalized by dividing them by the pre-injection average and compared between different treatment groups. **p<0.01 (saline; N = 8 mice, phlorizin; N = 7 mice, two-sample t-test) (D) Series of transcranial cortex-wide Ca^2+^ imaging under IG glucose injection with saline (control), raclopride (D2DR antagonist), and SCH23390 (D1DR antagonist) administered intraperitoneally. (E) Illustration of cortical activity patterns under IG glucose injection with raclopride (upper) and with SCH23390 (lower). The analysis focused on the earliest Ca^2+^ wave occurring within 3–8 seconds after injection. The pseudocolor representation uses the peak of the Ca^2+^ transient as the maximum value - 1SD and the mean + 1SD as the minimum value. (F) Comparative analysis of activation levels in each cortical region of mice under IG glucose injection with i.p. saline, raclopride, and SCH23390 injection. The methodology involved comparing the peak fluorescence intensity values of each brain region against those of a reference region (auditory cortex), covering the 50 seconds prior to injection and the subsequent post-injection period. Post-injection values were normalized by dividing them by the pre-injection average and compared across different treatment groups. *p<0.05 (saline, N = 7 mice, raclopride; N = 7 mice, SCH23390, N = 8 mice, one-way ANOVA followed by Tukey-Kramer method)

In the presence of phlorizin, the activation of the frontal cortex under IG glucose injection was not observed (**Figs. 3B-C**, Frontal: Saline vs. Phlorizin: 1.01 ± 0.20 vs. 0.91 ± 0.15, p = 0.0080, Somato: Saline vs. Phlorizin: 1.08 ± 0.17 vs. 1.00 ± 0.12, p = 0.33, Visual: Saline vs. Phlorizin: 1.34 ± 0.42 vs. 0.97 ± 0.20, p = 0.078, and RSC: Saline vs. Phlorizin: 1.16 ± 0.19 vs. 1.00 ± 0.13, p = 0.095). By this result, it is revealed that not only left vagus nerve but also frontal cortex immediate activation under IG glucose injection mediates glucose absorption of neuropod cells. Previous reports suggested that the centrifugal vagus nerves, synapsed by neuropod cells, project to the nucleus of the solitary tract and immediately activates VTA dopaminergic neurons.^22^ To investigate the potential involvement of dopamine in the IG glucose injection induced frontal cortex activation seen in Figure 2, we conducted similar transcranial cortex-wide Ca^2+^ imaging in the presence of D1 dopamine receptor (D1DR) antagonist SCH23390 and D2 dopamine receptor (D2DR) antagonist raclopride. The activation of the frontal cortex under IG glucose injection was absent in the presence of both the SCH23390 (D1DR antagonist) and the raclopride (D2DR antagonist) groups, compared to the control group (**Figs. 3D-F**, Frontal: Saline vs. Raclopride, 1.26 ± 0.25 vs. 0.94 ± 0.22, p = 0.014, Raclopride vs SCH23390, 0.94 ± 0.22 vs. 1.01 ± 0.07, p = 0.75, Saline vs SCH23390, 1.26 ± 0.25 vs. 1.01 ± 0.07, p = 0.05, Somato: Saline vs. Raclopride, 1.18 ± 0.29 vs. 0.96 ± 0.21, p = 0.16, Raclopride vs SCH23390, 0.96 ± 0.21 vs. 1.01 ± 0.09, p = 0.90, Saline vs SCH23390, 1.18 ± 0.29 vs. 1.01 ± 0.09, p = 0.29, Visual: Saline vs. Raclopride, 1.45 ± 0.58 vs. 1.06 ± 0.18, p = 0.13, Raclopride vs SCH23390, 1.06 ± 0.18 vs. 1.18 ± 0.17, p = 0.81, Saline vs SCH23390, 1.45 ± 0.58 vs. 1.18 ± 0.17, p = 0.32, and RSC: Saline vs. Raclopride, 1.38 ± 0.53 vs. 1.15 ± 0.29, p = 0.47, Raclopride vs SCH23390, 1.15 ± 0.29 vs. 1.18 ± 0.24, p = 0.99, Saline vs SCH23390, 1.38 ± 0.53 vs. 1.18 ± 0.14, p = 0.54). The results strongly suggest the involvement of neuropod cells in the immediate activation of the left vagus nerve and, furthermore, of the frontal cortex via central dopaminergic system.

### Immediate activation of the frontal cortex under IG glucose injection encompasses both neurons and astrocytes

The G7NG817 line transgenic mice used in Figures 2 and 3 revealed Ca^2+^ signaling from both neurons and astrocytes, rendering it ambiguous which cell type was implicated in the frontal cortex activation under IG glucose injection. To clarify this, cell-type-specific Ca^2+^ recording was performed on the frontal cortex using mice injected with AAV9 CaMKⅡa-GCaMP7f (targeting excitatory neurons) and Mlc1-tTA::tetO-YC-Nano50 transgenic mice (targeting astrocytes), accompanied by fiber photometry (**Figs. 4A-B**, **Supplementary Fig. 1**). The analysis indicated significant augmentation in fluorescence intensity within both neurons and astrocytes in the frontal cortex under IG glucose injection compared to water (**Figs. 4C-D**, Neurons: Water vs Glucose, 0.29 ± 0.67 vs. 1.85 ± 1.35, p = 0.049; Astrocytes: Water vs Glucose, 0.48 ± 0.28 vs. 1.46 ± 0.51, p = 0.006).

**Fig. 4.**
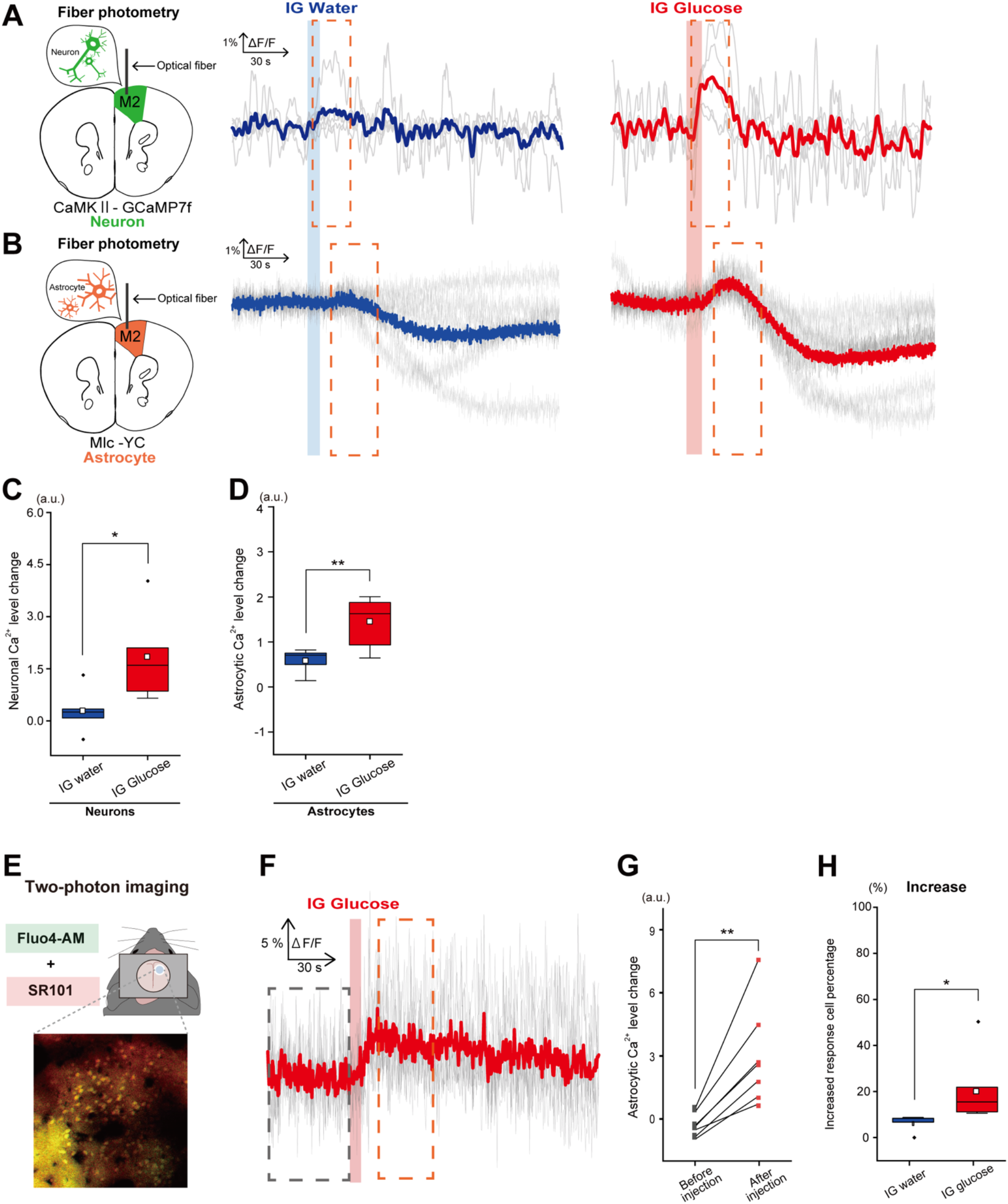
Neuronal and astrocytic Ca^2+^ dynamics in response to IG glucose and water injections. (A) Fiber photometric traces of averaged CaMKII-GCaMP7f (Neurons) signal responses to IG water injection (left) and IG glucose injection (right), with the light gray line depicting individual subject traces. (IG glucose; N = 5 mice, IG water; N = 5 mice) (B) Fiber photometric traces of averaged Mlc-YC (Astrocytes) signal responses to IG water injection (left) and IG glucose injection (right), with the light gray line representing individual subject traces. (IG glucose; N = 7 mice, IG water; N = 5 mice) (C) Comparison of Ca^2+^ activation levels under IG water (left) and glucose (right) injection of CaMKII-GCaMP7f (Neurons) mice. The analysis window corresponds to the area within the dotted line in (A). *p<0.05 (IG glucose; N = 5 mice, IG water; N = 5 mice, two-sample t-test) (D) Comparison of Ca^2+^ activation levels (early-phase) under IG water (left) and glucose (right) injection of Mlc-YC (Astrocytes) mice. The analysis window corresponds to the area within the dotted line in (B). **p<0.01 (IG glucose; N = 7 mice, IG water; N = 5 mice, two-sample t-test) (E) Representative two-photon Ca^2+^ imaging of layer 2 in the frontal cortex, with cells labeled with Fluo4-AM (green; Ca^2+^ indicator) and SR101 (red; astrocytes). (F) ΔF/F traces of fluorescence intensity for the cell population judged to exhibit an increase under IG glucose injection (gray line) alongside the corresponding average data (bold red line). (25 cells from N = 7 mice) (G) Comparison within each individual of the average value of astrocytic ΔF/F traces in response to IG glucose injection (15-38 seconds post-IG glucose injection; within the red dotted line in (G)) against the spontaneous state (50 seconds prior to injection; within the gray dotted line in (G)). **p<0.01 (IG glucose; N = 7 mice, two-sample repeated t-test) (H) Comparison of the percentage of SR101 positive cells exhibiting an Fluo4-AM signal increase among all SR101positive cells for each individual. *p<0.05 (IG glucose; N = 7 mice, IG water; N = 5 mice, two-sample t-test)

While IG glucose injection elicited activation in both neurons and astrocytes within the frontal cortex, the activation patterns were distinct between the two cell types. Neurons displayed a transient surge in fluorescence intensity, subsequently reverting to baseline, whereas astrocytes exhibited a pattern of where fluorescence intensity dipped below baseline after the transient increase. This reduction in fluorescence intensity occurred roughly 40 seconds post-injection of both glucose and water, with no significant difference between two (see discussion, **Supplementary Figs. 2A-B**).

The fluorescence signals captured via fiber photometry reflect Ca^2+^ activity at the population level among cell groups expressing Ca^2+^-sensitive fluorescent proteins within the analyzed brain region. The results of fiber photometry revealed that while neurons exhibit a simple Ca^2+^ response (increase or unincrease), astrocytes demonstrate complex response patterns that Ca^2+^ level decrease in the late phase after IG injection. To investigate whether such complex response was due to synchronous cellular activity triggered by some signal occurring in the late phase, we conducted two-photon Ca^2+^ imaging (**Fig. 4E**). We simultaneously addressed the possibility that the complex signal fluctuations observed only in astrocytes could be attributed to differences in the Kd values of the GCamp and YC probes by using Fluo-4 as a two-photon Ca^2+^ indicator. The findings revealed that Ca^2+^ level of some cells labeled with sulforhodamine 101 (SR101), a marker of astrocytes, increase under IG glucose injection (**Figs. 4F-G**, Before injection vs After injection, −0.12 ± 0.22 vs. 3.04 ± 2.39, p = 0.01). The increased cell percentages that fluorescence intensity change exceeding average + 0.5 × standard deviation (SD) under IG glucose injection was approximately 21%, and this proportion significantly greater than that observed under water injection (**Figs. 4H**, Water vs. Glucose, 6.01 ± 3.52 vs. 20.0 ± 13.4, p = 0.036). These results indicate the involvement of both neurons and astrocytes in the immediate activation of the frontal cortex under IG glucose injection.

### The activation of the frontal cortex under IG glucose injection is mediated by dopamine, with astrocytes and neurons responding through distinct receptors

Our transcranial Ca^2+^ imaging study revealed that the role of dopamine signal mediated by D1DR and D2DR for the activation of frontal cortex (**Figs. 3D-F**). Typically, frontal cortex neurons propagate reward-related signals via D1DR.^59–62^ Though cellular level astrocytes response to dopamine via each receptor is well investigated, it is not much investigated that the detail mechanism of astrocytes to dopamine in frontal cortex under physiological condition. ^63–66^ To delineate the contributions of D1DR and D2DR to the activation of neurons and astrocytes under IG glucose injection, the effect of D1DR and D2DR antagonist on CaMKII-GCamp7f and Mlc-YC signals were assessed using fiber photometry (**Fig. 5**).

**Fig. 5.**
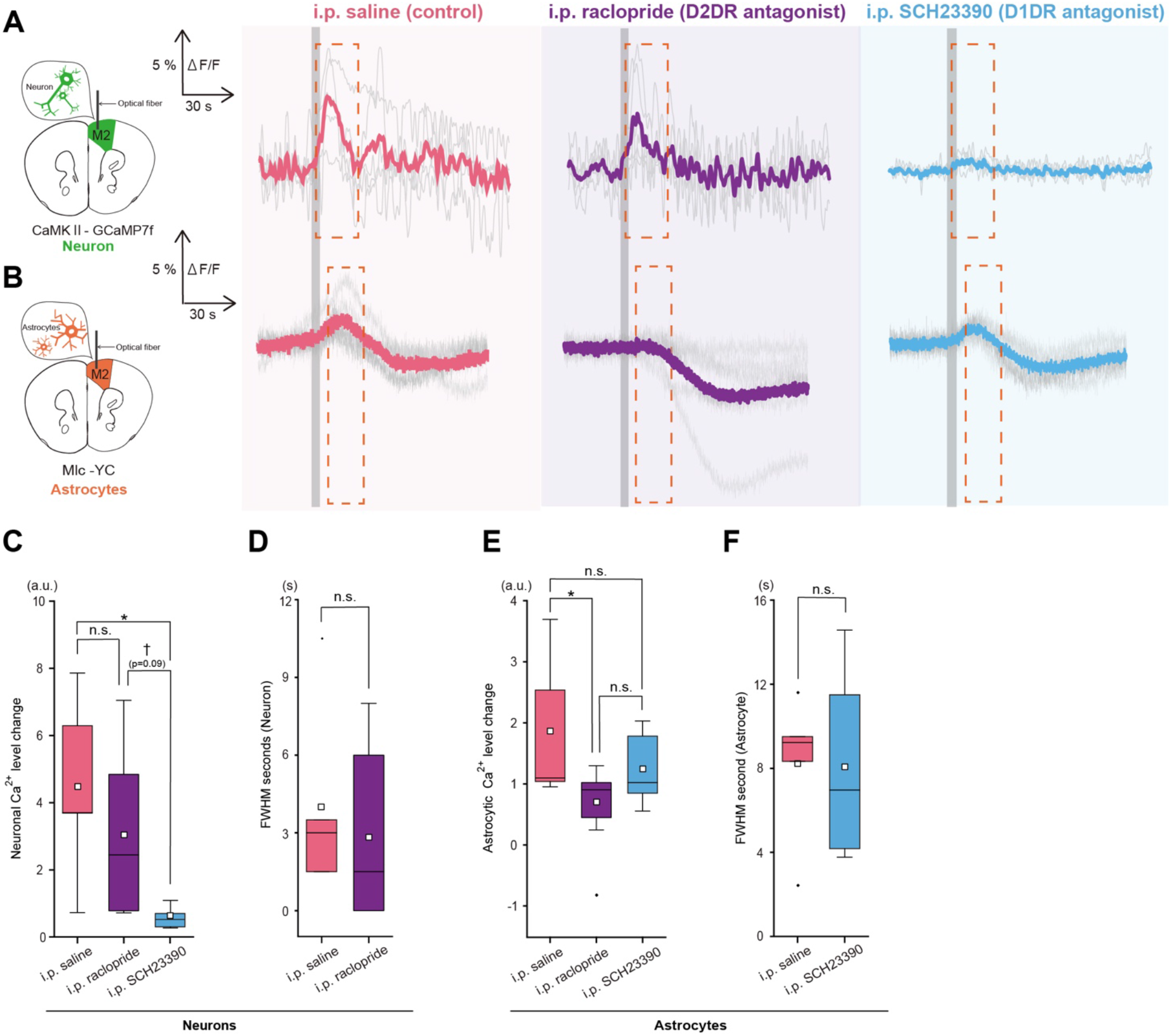
Differential responses of neuronal and astrocytic signals to IG glucose injection modulated by dopaminergic antagonists. (A) Fiber photometric traces of averaged CaMKII-GCaMP7f (Neurons) signal responses to IG glucose injection with saline (left), raclopride (middle), and SCH23390 (right) pretreatment. The gray shaded area denotes the IG glucose injection period. (saline; N = 5 mice, raclopride; N = 6 mice, SCH23390; N = 5 mice) (B) Fiber photometric traces of averaged Mlc-YC (Astrocytes) signal responses to IG glucose injection with saline (left), raclopride (middle), and SCH23390 (right) pretreatment. The gray shaded area indicates the IG glucose injection period. (saline; N = 5 mice, raclopride; N = 8 mice, SCH23390; N = 7 mice) (C) Comparison of Ca^2+^ activation levels under IG glucose injection of CaMKII-GCaMP7f (Neurons) mice with saline (left), raclopride (middle), and SCH23390 (right) pretreatment. †<0.1, *p<0.05 (saline; N = 5 mice, raclopride; N = 6 mice, SCH23390; N = 5 mice, one-way ANOVA followed by Tukey-Kramer method) (D) Comparison of the Half-Width at Half Maximum (HWHM) of CaMKII-GCaMP7f (Neurons) signals following IG glucose injection with saline (pink) and raclopride (purple) pretreatment. (saline; N = 5 mice, raclopride; N = 6 mice, t-test) (E) Comparison of Ca^2+^ activation (early-phase) levels under IG glucose injection of Mlc-YC (Astrocytes) mice with saline (left), raclopride (middle), and SCH23390 (right) pretreatment. *p<0.05 (saline; N = 5 mice, raclopride; N = 8 mice, SCH23390; N = 7 mice, one-way ANOVA followed by Tukey-Kramer method) (F) Comparison of the Half-Width at Half Maximum (HWHM) of Mlc-YC (Astrocytes) signals under IG glucose injection with saline (pink) and SCH23390 (light blue) pretreatment. (saline; N = 5 mice, SCH23390; N = 7 mice, t-test)

The findings indicated a significant reduction in the fluorescence intensity increase after IG glucose injection in neurons by the administration of D1DR antagonist (SCH23390), in contrast to the control group. However, a persistent elevation in fluorescence intensity was observed in both the control and D2DR antagonist (Raclopride) groups, with no significant differences in amplitude or activation duration between them (**Figs. 5A, C-D**, Amplitude: Saline vs. Raclopride, 4.45 ± 2.74 vs. 3.05 ± 2.08, p = 0.55, Raclopride vs SCH23390, 3.05 ± 2.08 vs. 0.58 ± 0.34, p = 0.18, Saline vs. SCH23390, 4.45 ± 2.74 vs. 0.58 ± 0.34, p = 0.036; HWHM: Saline vs. Raclopride, 3.74 ± 1.67 vs. 3.42 ± 1.69, p = 0.60).

In astrocytes, a significant reduction in fluorescence intensity increase was only noted in the group treated with the D2DR antagonist (raclopride), as opposed to the control group (**Supplementary Figs. 2C-D**). Conversely elevation in fluorescence intensity under IG glucose injection was still observed in both the control group and the group treated with the D1DR antagonist with no differences in amplitude or HWHM between these groups (**Figs. 5B, E-F**, Amplitude: Saline vs. Raclopride, 1.86 ± 1.21 vs. 0.60 ± 0.70, p = 0.044, Raclopride vs SCH23390, 0.60 ± 0.70 vs. 1.25 ± 0.55, p = 0.325, Saline vs. SCH23390, 1.86 ± 1.21 vs. 1.25 ± 0.55, p = 0.42; HWHM: Saline vs. SCH23390, 3.45 ± 1.54 vs. 4.11 ± 1.55, p = 0.95).

Taken together, these findings suggest that the activation of the frontal cortex under IG glucose injection could be comprised of neuronal and astrocytic activation through D1DR and D2DR-mediated dopaminergic signaling, respectively. Also, it probably facilitated by afferent vagus nerve activation via neuropod cells.

### Chronic mild restraint stress (CMRS) dampens immediate activation of the vagus nerve and frontal cortex under IG glucose injection

Finally, to investigate whether the immediate signals following glucose uptake, as revealed by previous experiment results, are involved in the decreased sucrose preference caused by chronic restraint stress, we examined the effects of mouse CMRS on activation of the left vagus nerve and frontal cortex. A chronic restraint stress model was established by confining mice in plastic tubes for 6.5 hours daily for 10 days, followed by conducting sucrose and glucose preference before and after stress exposure (**Fig. 6A**). The results demonstrated a significant decrease in preference in both tests (**Fig. 6B**, Sucrose preference test: Pre-stress vs. Post-stress, 0.87 ± 0.06 vs. 0.75 ± 0.08, p = 0.048; Glucose preference test, Pre-stress vs. Post-stress, 0.99 ± 0.02 vs. 0.86 ± 0.16, p = 0.059).

**Fig. 6.**
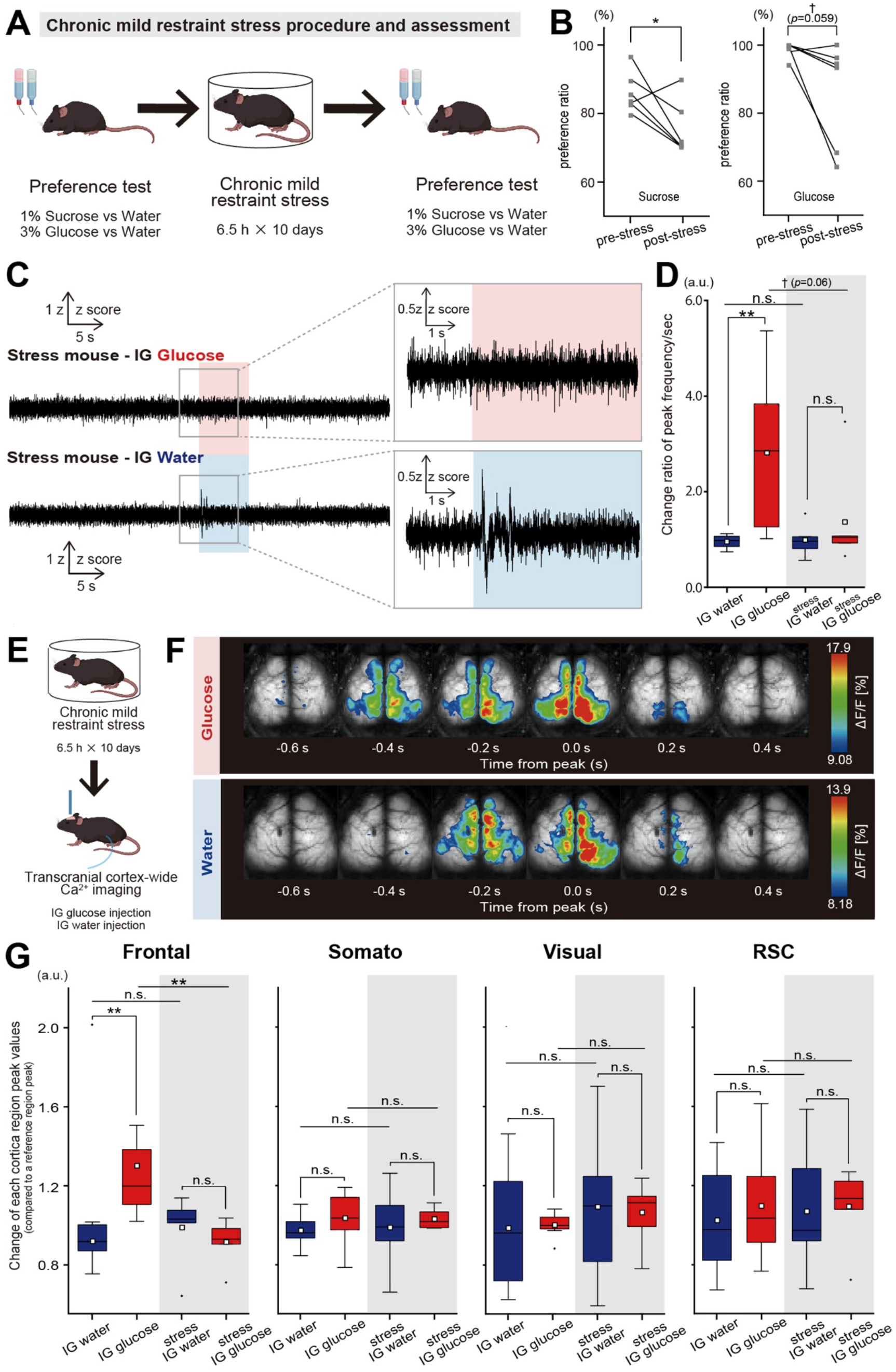
Impact of chronic mild restraint stress (CMRS) on vagus and cortical responses to IG glucose injection. (A) Overview of the CMRS methodology and assessment. (B) Changes in the preference ratios for 1% Sucrose (left) and 3% Glucose (right) pre-stress and post-stress. *p<0.05, †<0.1 (Sucrose preference test; N = 6 groups derived from 12 mice, two-sample paired t-test. Glucose preference test; N = 6 groups from 12 mice, Wilcoxon signed-rank sum test) (C) Illustration of normalized 50Hz high-pass filtered vagal nerve activity under IG glucose (upper) and water (lower) injection in CMRS-subjected mice. The periods of injection are highlighted in light red and light blue, respectively. (glucose: N = 6 mice, water: N = 5 mice) (D) Comparison of changes in the frequency of peaks per second in the vagus nerve pre- and post-IG injection between the water group (blue) and the glucose group (red). The data on a white background represents the group without CMRS, whereas the data on a gray background represents the group post-CMRS exposure. The initial 30 seconds before IG injection are considered the pre-stimulation period, and the 2-4 seconds after IG injection are considered the post-stimulation period. **p<0.01, †<0.1 (Untreated IG water: N = 8 mice, Untreated IG glucose: N = 9 mice, Stressed IG water: N = 5 mice, Stressed IG glucose: N = 6 mice, one-way ANOVA followed by Tukey-Kramer method) (E) Sequence of transcranial cortex-wide Ca^2+^ imaging post-CMRS. (F) Depiction of cortical activity patterns under IG glucose (upper) and water (lower) injection in CMRS-subjected mice. The target calcium wave for analysis is the earliest wave appearing within 3–8 seconds after injection. The pseudocolor representation employs the peak of the Ca^2+^ transient as the maximum value - 1SD and the mean + 1SD as the minimum value. Note that the frontal cortex activation under IG glucose injection was not observed in CMRS-subjected mice. (G) Comparison of activation levels in each cortical region of post-stress mice under IG water (blue) or glucose (red) injection. The analysis entailed comparing the peak fluorescence intensity values of each brain region against those of a reference region (auditory cortex) over a designated time window including the 50 seconds before injection and the post-injection period. Post-injection values were normalized by dividing them by the pre-injection average and compared between different treatment groups. The data on a white background represents the group without CMRS, whereas the data on a gray background represents the group post-CMRS exposure. **p<0.01 (Untreated IG water: N = 8 mice, Untreated IG glucose: N = 8 mice, Stressed IG water: N = 7 mice, Stressed IG glucose: N = 6 mice, one-way ANOVA followed by Tukey-Kramer method)

Subsequent measurements of vagus nerve activity under IG glucose/water injection in CMRS mice (**Fig. 6C**) revealed that the activation observed under glucose injection in healthy mice was absent in stress-subjected mice (**Fig. 6D**, Untreated: Water vs. Glucose, 0.96 ± 0.14 vs. 2.81 ± 1.56, p = 0.006, Stressed: Water vs. Glucose, 0.99 ± 0.36 vs. 1.34 ± 1.03, p = 0.93; Untreated water vs. Stressed water, 0.96 ± 0.14 vs. 0.99 ± 0.36, p = 0.99, Untreated glucose vs. Stressed glucose, 2.81 ± 1.56 vs. 1.34 ± 1.03, p = 0.06).

Given the reduced vagus nerve activation in CMRS mice, a corresponding decrease in frontal cortex activation, typically noted in healthy subjects, was anticipated. Transcranial cortex-wide Ca^2+^ imaging under IG glucose/water injection (**Figs. 6E-F**) confirmed a significant reduction in frontal cortex activation under IG glucose injection in CMRS mice, with no significant amplitude difference relative to water injection (**Fig. 6G**, Frontal: Untreated water vs. Untreated glucose, 0.92 ± 0.09 vs. 1.30 ± 0.32, p = 0.005, Stressed water vs. Stressed glucose, 0.99 ± 0.18 vs. 0.92 ± 0.11, p = 0.93, Untreated water vs. Stressed water, 0.92 ± 0.09 vs. 0.99 ± 0.18, p = 0.92, Untreated glucose vs. Stressed glucose, 1.30 ± 0.32 vs. 0.92 ± 0.11, p = 0.009; Somato: Untreated water vs. Untreated glucose, 0.97 ± 0.08 vs. 1.04 ± 0.13, p = 0.75, Stressed water vs. Stressed glucose, 0.99 ± 0.20 vs. 1.03 ± 0.05, p = 0.93, Untreated water vs. Stressed water, 0.97 ± 0.08 vs. 0.99 ± 0.20, p = 0.99, Untreated glucose vs. Stressed glucose, 1.04 ± 0.13 vs. 1.03 ± 0.05, p = 0.99; Visual: Untreated water vs. Untreated glucose, 0.99 ± 0.30 vs. 1.00 ± 0.06, p = 0.99, Stressed water vs. Stressed glucose, 1.09 ± 0.39 vs. 1.07 ± 0.16, p = 0.99, Untreated water vs. Stressed water, 0.99 ± 0.30 vs. 1.09 ± 0.39, p = 0.86, Untreated glucose vs. Stressed glucose, 1.00 ± 0.06 vs. 1.07 ± 0.16, p = 0.97; RSC: Untreated water vs. Untreated glucose, 1.02 ± 0.27 vs. 1.1 ± 0.27, p = 0.95, Stressed water vs. Stressed glucose, 1.07 ± 0.32 vs. 1.09 ± 0.20, p = 0.99, Untreated water vs. Stressed water, 1.02 ± 0.27 vs. 1.07 ± 0.32, p = 0.99, Untreated glucose vs. Stressed glucose, 1.1 ± 0.27 vs. 1.09 ± 0.20, p = 0.99). These results suggest disruption in the immediate vagus nerve response under gut glucose uptake in CMRS mice.

## Discussion

In this study, we elucidated that the activation of the vagus nerve induced by direct duodenal glucose injection (IG glucose injection) is mediated by neuropod cells via SGLT1 (**Fig. 1**). Furthermore, we demonstrated that this signal activates both astrocytes and neurons in the frontal cortex through the application of transcranial cortex-wide Ca^2+^ imaging (**Figs. 2 and 3**) and fiber photometry (**Figs. 4 and 5**). Pharmacological evidence confirmed the involvement of D2DR in astrocytic activation and D1DR in neuronal activation (**Figs. 3 and 5**). Additionally, we identified alterations in signals originating from neuropod cells in mice subjected to chronic mild restraint stress (CMRS) at the level of vagus nerve activation (**Fig. 6**).

### The immediate activation of the frontal cortex following intestinal glucose administration would be attributed to the activation of the Mesocortical dopaminergic pathway derived from neuroepithelial cells

Previous research has shown that sugar absorption in the gut triggers the activation of central dopaminergic neurons. These dopaminergic pathways can be broadly divided into four: the nigrostriatal pathway, tuberoinfundibular pathway, mesocortical pathway and mesolimbic pathway. Among those four pathways, it has been shown that IG sugar injection can activate nigrostriatal and mesolimbic dopaminergic pathways some minutes after the start of the IG sugar injection.^2,3,7,22–27^ Early research focused on dorsal and ventral striatum where nigrostriatal and mesolimbic dopaminergic pathways’ projection target, probably because the experimental techniques used, such as microdialysis and virus tracing, are more suited to detecting long-lasting dopaminergic reactions in the dorsal pathway.^7,23,26^

On the other hand, recent research revealed that an IG sugar injection activates the VTA—a component of the mesolimbic and mesocortical pathway immediately, occurring within seconds, compared to the timing of dopamine detection in the striatum. This activation is expected to be initiated by the neuropod cell that synapses with the left vagus nerve.^22^ Our results demonstrated that an IG glucose injection immediately activates the frontal cortex through a central dopaminergic pathway originating from neuropod cells, as illustrated in **Figs. 3A-F**. There is a possibility that the mesocortical pathway, which originates from the VTA and shares similarities with the mesolimbic pathway, may mediate the immediate activation of the frontal cortex derived from neuropod cells. This is supported by our experimental results, where the activation of the frontal cortex induced by IG glucose injection was inhibited by dopamine receptor antagonists.^39^

This is significant because dopamine-induced activation in the frontal cortex plays an important role in reward-based decision-making. Our results provide essential insights into understanding how signals from neuropod cells may enhance sugar preference.

### The physiological significance of the immediate activation of both astrocytes and neurons in the frontal cortex by dopamine under IG glucose injection

Using fiber photometry, we confirmed that both astrocytes and neurons in frontal cortex activate immediately under IG glucose injection. Furthermore, our pharmacological experiment revealed that the increase in Ca^2+^ levels in astrocytic and neuronal induced by dopamine occurred via D2DR and D1DR, respectively. It is known that astrocytes release ATP/adenosine as gliotransmitters upon activation by dopamine through D2DR. These molecules, in turn, activate neuronal adenosine 1 receptors (A1 receptors), leading to a reduction in neuronal activity.^63–66^ Interestingly, the disruption of this astrocyte-dependent dopamine regulation in the frontal cortex is linked to the development of obsessive-compulsive spectrum disorders (OCD) because of dysregulation of the corticostriatal circuit.^44^ The corticostriatal circuit, which involves projection from frontal cortex neurons to the striatum, plays an important role in food-seeking behavior.^43^

Although our experiment did not directly validate the release of ATP/adenosine following astrocytic activation via D2DR and the subsequent suppression of neuronal activity, the observed increase in intracellular Ca^2+^ concentration in astrocytes following dopamine reception via D2DR could be considered a key factor in modulating sugar intake in mice.

### Astrocytes in frontal cortex response to dopamine in different patterns

Our fiber photometric recording following IG glucose injection revealed though neurons exhibited a transient increase in fluorescence intensity following the IG glucose injection, astrocytes exhibited a complex response patterns to dopamine in the frontal cortex (**Fig. 4B**). Specifically, we observed an initial transient increase in fluorescence intensity, followed by a reduction approximately 40 seconds post-injection. This decline was also noted in the groups receiving IG water injections and those treated with dopamine antagonists post-IG glucose injection (**Supplementary Figs. 2A-D**). These observations led us to hypothesize that the reduction in astrocytic Ca^2+^ levels during the latter phase might be due to mechanisms unrelated to the dopamine response induced by IG glucose injection.

To further investigate these response patterns, we examined the Ca^2+^ activity of individual astrocytes under IG glucose injection using two-photon imaging. We discovered diverse response patterns among frontal cortex astrocytes, indicating that the composite response observed in fiber photometry does not reflect synchronized activity across all response cells (**Figs. 4F-H and Supplementary Fig. 3C**). The responses were primarily categorized into three types: [1] cells with an increase in Ca^2+^, [2] cells with a decrease in Ca^2+^ and [3] cells showing no response. While a slightly higher proportion of cells exhibited a Ca^2+^ decrease in the glucose group, the difference was not statistically significant (p = 0.06). Conversely, the IG water injection group showed a significantly higher proportion of cells with no change in fluorescence intensity in response to stimulation (**Supplementary Figs. 3A-C**). At the cellular level, the onset of astrocytic Ca^2+^ decrease following IG injection was delayed by approximately 9 seconds compared to the Ca^2+^ elevation, with no significant differences in the timing of each response pattern (**Supplementary Figs. 3D-F**).

We speculate that the increase in fluorescence intensity in some cells may arise from the heterogeneity within the astrocyte population, a notion supported by observations in cultured astrocytes from prior research.^66^ As for the decrease in Ca^2+^ levels, it is expected that this result is not due to metabolite-derived signals during the latter phase post-IG injection but rather from other acute responses induced by the IG injection itself. The occurrence of intracellular Ca^2+^ decrease in astrocytes presents a compelling phenomenon, with many aspects yet to be fully understood. Considering the diverse responses observed, such as the immediate decrease in heart rate and alterations in cerebrospinal fluid (CSF) upon artificial activation of vagus nerve fibers, as well as contradictory findings like the activation or inhibition of neurons in the frontal cortex, further exploration is necessary to identify the factors responsible for the acute decrease in astrocytic Ca^2+^ induced by the stimulation methodology applied in this study.^67–69^

### Chronic mild restraint stress (CMRS) decreases the immediate activation of left vagus nerve under IG glucose injection

Chronic stress has been demonstrated to reduce sucrose preference in mice,^30,70^ suggesting alterations in reward processing mechanisms. Such chronic stress is known to induce various changes within the brain, particularly affecting dopamine neurotransmission^46–48,50,52^, and also impacts the activity of the autonomic nervous system, including vagus nerve function. The specific effect of stress on post-stress vagus nerve activity, including deviations in spike frequency bands and aberrant vagus nerve signals, are complex and warrant further investigation, as they may contribute to brain alterations.^49,51,71^

Our results indicated no significant difference in the peak response of the left vagus nerve to IG glucose injection between the stressed group receiving IG water and those receiving IG glucose. Furthermore, there was no significant difference in the IG water injection responses between stressed and healthy mice, but a modest and significant difference was noted in the IG glucose injection response between stressed and control mice (**Figs. 6C-D**, IG glucose injection + CMRS vs IG glucose injection, 2.81 ± 1.56 vs. 1.34 ± 1.03, p = 0.06). Based on these observations, we hypothesized that CMRS-exposed mice might exhibit reduced activation of the frontal cortex in response to IG glucose injection. This hypothesis was confirmed by observing a significant reduction in frontal cortex activation in CMRS-exposed mice in response to IG glucose injection, with no significant deviation from intragastric water injection responses (**Figs. 6F-G**).

The role of dopamine-induced frontal cortex activation in reward-based decision-making is well-established.^39–43^ Our results suggest that the decreased sucrose preference in CMRS-exposed mice could be partially attributed to reduced activation of the left vagus nerve, possibly originating from neuroepithelial cells. However, our experiments did not verify whether the decrease in sucrose preference could be ameliorated by specifically activating neuroepithelial cells in post-CMRS-exposed mice. Given previous research that has shown specific activation of neuroepithelial cell-derived signals during a two-bottle choice test of sucrose (including glucose) versus sucralose (not including glucose) leads to a stronger preference for glucose-containing sucrose in comparison to control,^34^ further studies are needed to directly investigate the contribution of neuroepithelial cell-derived signals to the reduced sucrose preference observed in post-CMRS-exposed mice.

Our study demonstrated that intragastric (duodenum) glucose injection immediately activates the left vagus nerve and frontal cortex within seconds, likely mediated by neuropod cells that synapse with the vagus nerve almost instantaneously. Additionally, we observed that frontal cortex activation levels immediately following IG glucose injection are diminished under CMRS conditions, which may be due to vagus nerve activity abnormalities induced by CMRS.

The activation of the frontal cortex by dopamine plays an important role in reward-based decision-making and regulating striatal activity by modulating corticostriatal circuit connectivity.^39–44^ While our research concentrated on the ventral dopaminergic pathway, many research have focused on the dorsal striatum following dorsal dopaminergic pathway activation and have shown that dopamine uptake in dorsal striatum post-gastric glucose absorption stimulates sugar intake in mice.^18,23,26,32,43^

We have yet determined whether frontal cortex activation directly enhances sucrose intake in mice or if changes in the activity of other brain regions, such as the striatum, induced by frontal cortex activation, indirectly facilitate sucrose intake. Nonetheless, our results suggest that the immediate activation of the frontal cortex following glucose absorption in the gut is crucial for sucrose preference in mice. Furthermore, any abnormalities in this activation process may contribute to the reduced sucrose preference observed in mice subjected to chronic stress.

## Methods

All experimental protocols were approved by the Institutional Animal Care and Use Committee of Ochanomizu University, Japan (animal study protocols 23006). All animal experiments were performed according to the guidelines for animal experimentation of Ochanomizu University that conforms with the Fundamental Guidelines for Proper Conduct of Animal Experiment and Related Activities in Academic Research Institutions (Ministry of Education, Culture, Sports, Science and Technology, Japan). Efforts were taken to minimize the number of animals used. This study was carried out in compliance with the ARRIVE guidelines.

### Animals

Adult male and female wildtype, G7NG817 transgenic, and Mlc-YC (Mlc-tTA::tetO-yellow Cameleon-Nano50) double transgenic mice (older than 8 weeks) were used. ^54,72^ The background strain of all mice is C57BL/6. Mice were housed under a 12 h /12 h light/dark cycle and raised in groups of up to five mice. G7NG817 mice was obtained from the RIKEN Bio Resource Center (Resource IDs: RBRC09650).

### Catheter insertion for proximal duodenum

Under 2.0% isoflurane anesthesia, the catheter insertion procedure was conducted as previously described.^73^ Mice were placed on a surgical table with a heat pad maintained at 37℃. A skin incision of approximately 1.5 cm was made 1 cm to the right of the abdominal median and 5 mm below the xiphoid process. Then, a 1.5 cm incision was created in the abdominal wall at the same location as the skin incision. A small perforation (approximately 1.5mm in diameter) was made in the pyloric antrum, and the catheter tip was inserted. The catheter was secured, and the abdominal wall sutured, allowing the catheter to protrude externally, using a 5/0 silk suture. The skin incision was then sutured in a similar manner to the abdominal closure. The catheter was flushed with saline and sealed with a needle cap to prevent bacterial infection. Mice were then returned to their cage for a minimum recovery period of 48 hours.

### Preparation for *in vivo* transcranial Ca^2+^ imaging

Under 2.0% isoflurane anesthesia, preparation was performed a day prior to the imaging session as previously outlined.^73^ The mouse was positioned on a stereotaxic platform, and the scalp hair was removed using hair removal cream. After applying local anesthetic gel, the scalp was excised completely. The periosteum’s connective tissue was removed, and acrylic cement was promptly applied to the skull to prevent opacity from evaporation. The skull surface was allowed to dry for 5 minutes.

### Optical fiber implantation

Stereotaxic surgery was performed under anesthesia with a ketamine-xylazine mixture (100 and 10 mg/kg, i.p.). For fiber photometric recordings, an optical fiber cannula (CFMC14L05, ⌀ 400 mm, 0.39 NA, Thorlabs) was implanted into layer 5 of the right secondary motor cortex (M2) in one direction (anteroposterior (AP), +1.8 mm; Mediolateral from Bregma (BL), 0.7 mm; dorsoventral from the skull surface (DV), 1.4 mm).

### Virus injection

For microinjection of AAV into the brain for fiber photometric recording, AAV9-CaMKIIa-jGCaMP7f (3.0 × 10^13^ vg per mL)^74^ was injected into the layer5 right secondary motor cortex (AP, +1.8 mm, ML, 0.7 mm, DV, 1.4 mm). AAV microinjection was performed using a stainless-steel microinjection cannula (CXMI-4T, Eicom) attached with a 10-mL Hamilton syringe directed by a syringe pump (Legato130, KD Scientific) at a flow rate of 0.04 mL min^−1^ for 10 min. AAV microinjection was carried out 3 weeks before surgery for fiber implantation.

### Surgery procedures for two-photon imaging

For two-photon imaging, a craniotomy with a diameter of 2 mm was made above the frontal cortex (AP, +1.65 mm, ML, 1.35 mm). The dura mater was surgically removed. Fluo4-AM (Thermo Fisher Scientific, 1μM, dissolved in a DMSO stock solution containing 10% Pluronic F127) was topically applied to 2 hours to facilitate Ca^2+^ activity detection, followed by Sulforhodamine 101 (100 μM in saline) for 1 minute to specifically label astrocytes, which was then rinsed with HEPES-buffered artificial cerebrospinal fluid (aCSF). After the dye loading, the craniotomy site was covered with agarose (2.0% w/v in aCSF) and gently sealed with a thin glass coverslip (2.7 × 2.7 mm, thickness: 0.3 mm, Matsunami Glass). The cranial window was then secured to the skull using dental cement.

### Surgical procedure for cervical vagus nerve electrophysiological recording

Cuffs electrodes covered with a silicon tube (outer diameter: 0.7 mm; inner diameter: 0.3 mm, Unique Medical), connected to gold pins or plastic connectors, were attached to the left cervical vagus nerve. Under 2.0% isoflurane anesthesia, mice with previously attached catheter to the gut were secured on a surgical table equipped with a heating pad maintained at 37℃. The neck hair was removed by using a hair removal cream, then mice were applied local anesthetic gel. The ventral cervical area was incised by scissors, then salivary gland was cautiously removed by tweezers. The left vagus nerve was carefully dissected from the carotid sheath, and the cuff portion was attached to the nerve.

### Behavioral test

#### Sucrose vs. water and glucose vs. water preference test

Before chronic mild restraint stress (CMRS), pairs of mice were systematically paired while ensuring gender segregation. These pairs were then introduced into cages containing two bottles, designed to mitigate spillage through vibrations. To familiarize the mice with the presence of two bottles, both containers were initially filled solely with water for several days. One day prior to start the preference test, one bottle was filled with a sugar solution (1% sucrose solution for the sucrose preference test or 3% glucose solution for the glucose preference test), while the other retained water. Mice were granted unrestricted access to both bottles. On the day of the test, the weights of each bottle were measured, and their positions within the cage were alternated from the previous day. One day into the preference test, the positions of the bottles were swapped to counteract any side preference bias in the mice. Two days into the preference test, the weights of both bottles were recorded. One day after CMRS procedure, bottles containing a sugar solution (1% sucrose solution for the sucrose preference test or 3% glucose solution for the glucose preference test) and water were positioned in the cages where the stress procedure had been conducted. Mice were given free access to both bottles, and following the protocol established in the pre-stress test, the positions of the bottles switched one day after the test began. Two days after the test commenced, the bottles were collected, and their weights were measured. The preference for the 1% sucrose or 3% glucose solution was calculated as follows.

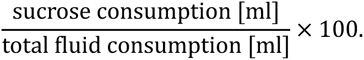

#### Chronic mild restraint stress (CMRS) procedure

Mice were paired with same sex counterparts and placed together in a small cage for 8hours a day for 10 consecutive days. The small cage is a plastic circle container (diameter; 7 cm, depth; 8 cm) which has several small holes to allow for ventilation of the interior air. Through the CMRS, each mouse had restricted to access water, but their food intake was unrestricted as pellet was placed in the plastic circle container.

### Drug application

The SGLT1 inhibitor, phlorizin, was dissolved in saline (100mM, MedChemExpress) and intraperitoneally injected at a dosage of 100μg/kg 20 minutes prior to initiating transcranial cortex-wide Ca^2+^ imaging or vagus nerve electrophysiological recordings. The D1DR antagonist SCH 23390 (0.25mg/kg, Sigma) was dissolved in saline and administered intraperitoneally 30 minutes before commencing transcranial cortex-wide Ca^2+^ imaging and fiber photometric recordings. The D2DR antagonist raclopride (400μg/kg, Sigma) was dissolved in saline or administered intraperitoneally 20 minutes before starting transcranial cortex-wide Ca^2+^ imaging or fiber photometric recordings.

### In vivo transcranial cortex-wide Ca^2+^ imaging

G7NG817 mice under 1.0% isoflurane anesthesia were placed onto stereotaxic platform using auxiliary ear bars under a fluorescence stereo microscope (MVX10, Evident). The U-MGFPHQ filter set (excitation 460-480 nm, emission 495-540 nm, Evident) was used with the U-HGLGPS light source (Evident). Images were acquired using the ORCA-spark CMOS camera (Hamamatsu Photonics) using HC Image software (Hamamatsu Photonics). Images were acquired with a size of 512 × 512 pixels and 16-bit resolution. ALL images were acquired at 10 Hz.

### In vivo two-photon imaging

Two-photon imaging was performed on 1.0% isoflurane-anaesthetized wiled type mice using a multi-photon laser scanning microscope (FVMPE-RS, Evident) equipped with an InSight laser system (Spectra-Physics) and an Olympus objective (FV30-AC25W, numerical aperture: 1.05, working distance: 2 mm, and immersion medium: water). Images through IG injection session were acquired using resonant-scanner at 512×512 pixels (16-bit depth) with a frame rate at 10Hz, whereas a single image before the IG injection session was acquired using a galvano-scanner at the same resolution to identify cells where merged green and red signals. The excitation wavelength was adjusted from 690 to 1100 nm. Laser emission wavelengths of 495-540 nm was used for Fluo4-AM and 575-645 nm was used for SR101 excitation, employing the FV30-FGR filter (Evident). Images were captured using Evident software at a frame rate of 10 Hz.

### Fiber photometory

To detect the Mlc-YC and CaMKII-GCamp fluorescence signals, a custom-made fiber photometric system designed by Olympus Engineering was used.^75^ For recording of Mlc-YC signal, input light (center wavelength in 435 nm; silver-LED-435, Prizmatix) was reflected off a dichroic mirror (DM455CFP, Olympus), coupled into an optical fiber (M41L01, ⌀ 600 mm, 0.48 NA, Thorlabs), linked next to an optical fiber (M79L01, ⌀ 400 mm, 0.39 NA, Thorlabs) through a pinhole (⌀ 600 mm). It was then delivered to an optical fiber cannula (CFMC14L05, Thorlabs) implanted into the mouse brain. The LED power was < 200 µW at the fiber tip. Emitted yellow and cyan fluorescence light from the YC probe was collected via an optical fiber cannula, divided by a dichroic mirror (DM515YFP, Olympus) into cyan (483/32 nm band-pass filter, Semrock) and yellow fluorescence (542/27 nm band-pass filter, Semrock), and detected using two distinct photomultiplier tubes (H7422-40, Hamamatsu Photonics). For recording of CaMKII-GCaMP signal, input light (center wavelength of 475 nm; silver-LED-475, Prizmatix) and dichroic mirrors (DM490GFP, Olympus and FF552-Di02–25×36, Semrock) were used to detect green fluorescence with a band-pass filter (BA495-540GFP, Olympus). For recording of the GCaMP isosbestic control signal, input light (center wavelength of 405 nm; M405F1, Thorlabs) and dichroic mirrors (DMLP425R, Thorlabs; DM490GFP, Olympus; and FF552-Di02–25×36, Semrock) were used to detect green fluorescence with a band-pass filter (BA495-540GFP, Olympus). Fluorescent signals were digitized using a data acquisition module (NI USB-6008, National Instruments) and recorded using a custom-made LabVIEW program (National Instruments). CaMKII-GCaMP and Mlc-YC signal was collected at 1 Hz and 1 kHz, respectively.

### Vagus nerve electrophysiological recording

Vagus nerve electrophysiological recording was performed in mice under 1.0% isoflurane anesthesia using a heating pad maintained at 37℃, with a cuff electrode and a signal amplifier (HAS-4 Head Amplifier System, Bio Research Center Co. Ltd.). Data were amplified via head stage (×1000, filtered 10–7000, Hz) and subsequently digitized using LabVIEW (National Instruments).

### Image processing

For transcranial images, images are binned 64 × 64 pixels, and hand-drawn regions of interest (ROIs) were determined using ImageJ. ROIs are aligned with the mouse brain atlas to designate ROIs as the frontal area (Frontal), the somatosensory area including the barrel area (Somato), the occipital area (Visual), and the retrosplenial region (RSC). ROIs coordinates were extracted using the MATLAB function ‘ReadImageJROI’. The averaged fluorescence intensity change rate (ΔF/F) within each ROI was computed using MATLAB, where ΔF/F is defined as follows.

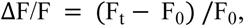

where F_t_ represents the fluorescence intensity value at a specific time and F_0_represents the average intensity value during the spontaneous state, defined as the 50 seconds preceding the injection start. The averaged fluorescence intensity changes rate of each ROI underwent a low-pass filter to restrict frequencies below 1Hz, and Ca^2+^ oscillation peaks exceeding the baseline intensity value of each ROI were identified by MATLAB. Peak ratios for Frontal, Somato, Visual, and RSC were determined by dividing each peak values by the peak value of the reference ROI (i.e. auditory area). Spontaneous peak ratio was the average over 50 seconds before injection start, and target waveforms were identified between 3 to 8 seconds after injection initiation. Activation levels were defined as the ratio of the under injection peak ratio to the spontaneous peak ratio.

For two-photon imaging analysis, ROIs were determined by manually outlining the merged areas both SR101 and Fluo4-AM positive cells. ROIs coordinates were extracted by using the MATLAB function ‘ReadImageJROI’. The averaged fluorescence intensity change rate (ΔF/F) within each ROI was computed using MATLAB, with ΔF/F defined as above. The average and standard deviation (SD) during the spontaneous state were computed. ROIs exhibiting fluorescence intensity exceeded <the spontaneous average + 0.5SD> within 15-38 seconds post-injection were defined as ‘Increase’ (this period is defined as the average duration for the increase in populational astrocytic Ca^2+^ response in the frontal cortex). ROIs with fluorescence intensity below <the spontaneous average - 0.5SD> during the same timeframe were defined as ‘Decrease’. ROIs not meeting either criterion were labeled as ‘Nonresponse’. The proportion of Increase, Decrease and Nonresponse ROIs for each individual was determined based on the count of each ROI type relative to the total number of ROIs per individual (Fig. 4G and Supplementary Fig. 3B). The average waveform of classified ROIs (Increase or Decrease) for individuals in the IG glucose injection group was calculated and using the waveforms, the peak onset, half-width at half maximum (HWHM), and peak time were computed for each individual (Supplementary Figs. 3D-F).

#### Fiber photometric imaging analysis

For neuronal and astrocytic Ca^2+^ upregulation analysis after IG injection, individual’s ΔF/F was calculated. ΔF/F was defined as follows.

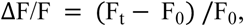

where F_t_ represents the fluorescence intensity value at a specific time and F_0_ represents the average intensity value of spontaneous state. Spontaneous state is defined as 30 seconds leading up to the start of the injection. In the data from individuals in the group subjected to IG glucose injection, the average and standard deviation (SD) during the spontaneous state were calculated. The upregulation time post-injection, where the fluorescence intensity exceeded <the average + 3 SD>, was determined for each individual animal. The analysis window for post-injection fluorescence intensity was defined as the time between the minimum and maximum values of the determined time of entire data. The average waveform of individual from the IG glucose injection group was calculated. Using the waveforms, the peak onset, half width at half maximum (HWHM) were computed. In the neuronal dataset, the difference between the median fluorescence intensity within the analysis window and the median fluorescence intensity during the spontaneous state was defined as the neuronal Ca^2+^ level upregulation change (Figs. 4C and 5C). In the astrocyte dataset, the difference between the top 30% of the average fluorescence intensity within the analysis window and the average fluorescence intensity during the spontaneous state was defined as the astrocytic Ca^2+^ level upregulation change (Figs. 4F and 5E).

For astrocytic Ca^2+^ downregulation analysis post-IG injection, the downregulation time post-injection, where the fluorescence intensity below <THE average - 3SD>, was determined for each individual animal. The analysis window for post-injection fluorescence intensity was defined as the time between the minimum and maximum values of the determined time of entire data. The difference between the average fluorescence intensity within the analysis window and the bottom 30% of the average fluorescence intensity during the spontaneous state was defined as the astrocytic Ca^2+^ level downregulation change (Supplementary Figs. 2B and 2D).

For vagus nerve electrophysiological recording analysis, recording data were subjected to high-pass filter, limiting them to frequencies above 50Hz. The filtered recording data’s average, maximum value and standard deviation in spontaneous state (30 seconds leading up to the start of the injection) was calculated, then the entire recording data was normalized with reference to the average and maximum values during the spontaneous state. The median and standard deviation of the normalized data during the spontaneous state were calculated. Peaks that exceeding (median + 2SD) were identified within the entire set of normalized data for each individual. Peaks surpassing the median + 5SD were removed as noise. Number of peaks were calculated. The number of peaks per second was determined, and a comparison was made between the average number of peaks during the spontaneous state and of peaks during the 2-4 seconds post-injection period.

## Acknowledgments

This work was supported by Ochanomizu University, the RIKEN CBS-EVIDENT Open Collaboration Center (BOCC), KAKENHI grants (18K14859, 20K15895), the JST FOREST Program (Grant Number JPMJFR204G), the Research Foundation for Opto-Science and Technology, the Kao Research Council for the Study of Healthcare Science, the Japan Association for Chemical Innovation, and the TERUMO LIFE SCIENCE FOUNDATION. We extend our gratitude to Takashi Shichita of Tokyo Medical and Dental University for generously providing AAV9-CaMKIIa-jGCaMP7f, and to Atsushi Miyawaki from the RIKEN Center for Brain Science for his expert supervision of the two-photon imaging. We also acknowledge that some of the figures were created using BioRender (https://www.biorender.com/).

---

**Supplementary Fig. 1.**
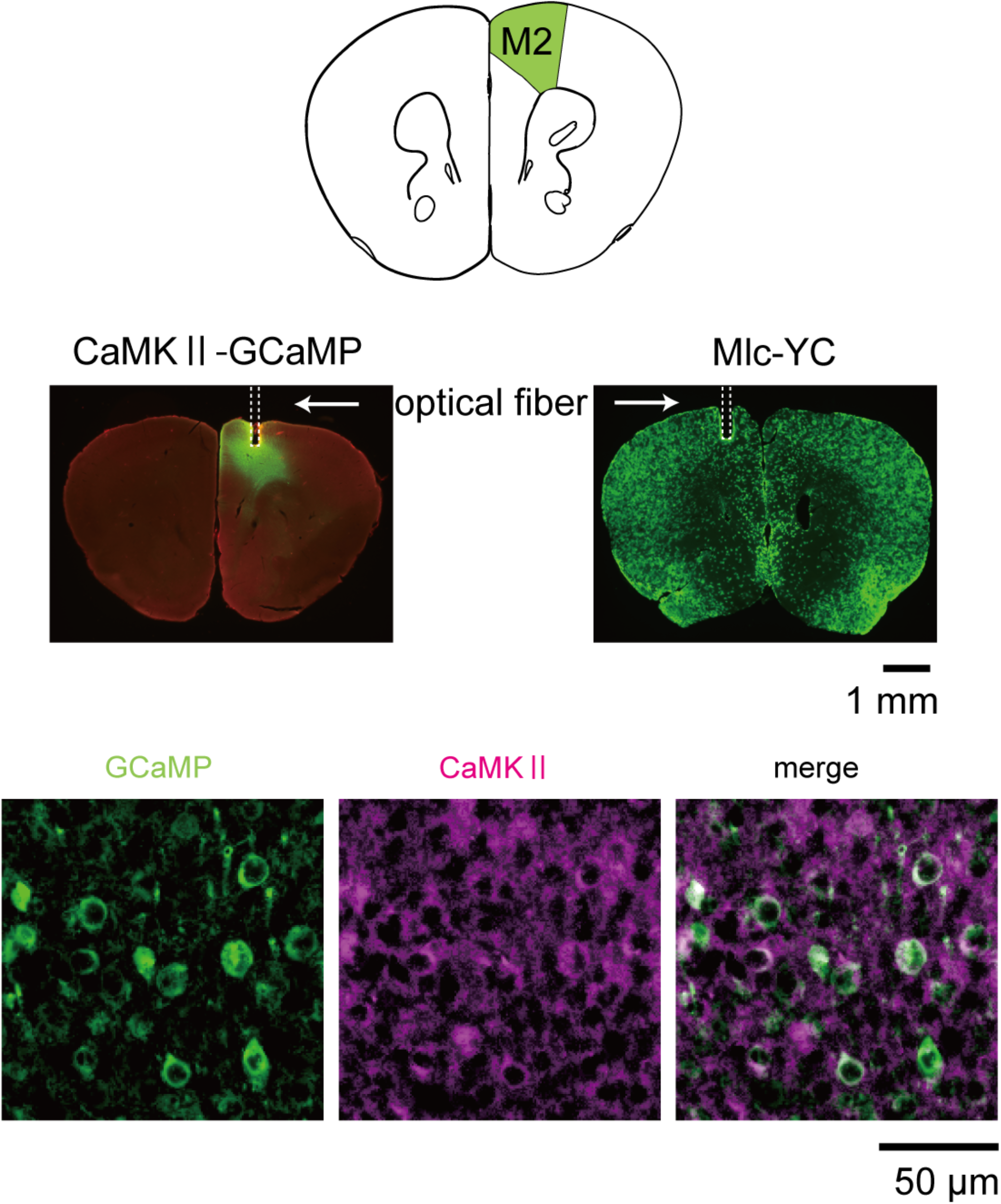
Histological Illustrations of CaMKII-GCaMP and Mlc-YC-nano50 probes. (top) Histological reconstructions depicting the placement of optical fiber tips in the M2 of CaMKII-GCaMP (left) and Mlc-YC (right) probes with green indicating GFP staining. Scale bar: 1 mm. In CaMKII-GCaMP probes, GFP immunostaining reveals CaMKII expression within pyramidal neurons. In Mlc-YC mice, GFP immunostaining indicates the presence of Yellow Cameleon Nano-50 driven by astrocyte-specific tetracycline transactivator expression. (bottom) GFP fluorescence microscopy in layer 5 of the secondary frontal cortex (M2) showing GCaMP (left), CaMKII (middle), and their composite image (right): Green denotes GCaMP fluorescence, while pink highlights CaMKII labeling. Scale bar: 50 μm.

**Supplementary Fig. 2.**
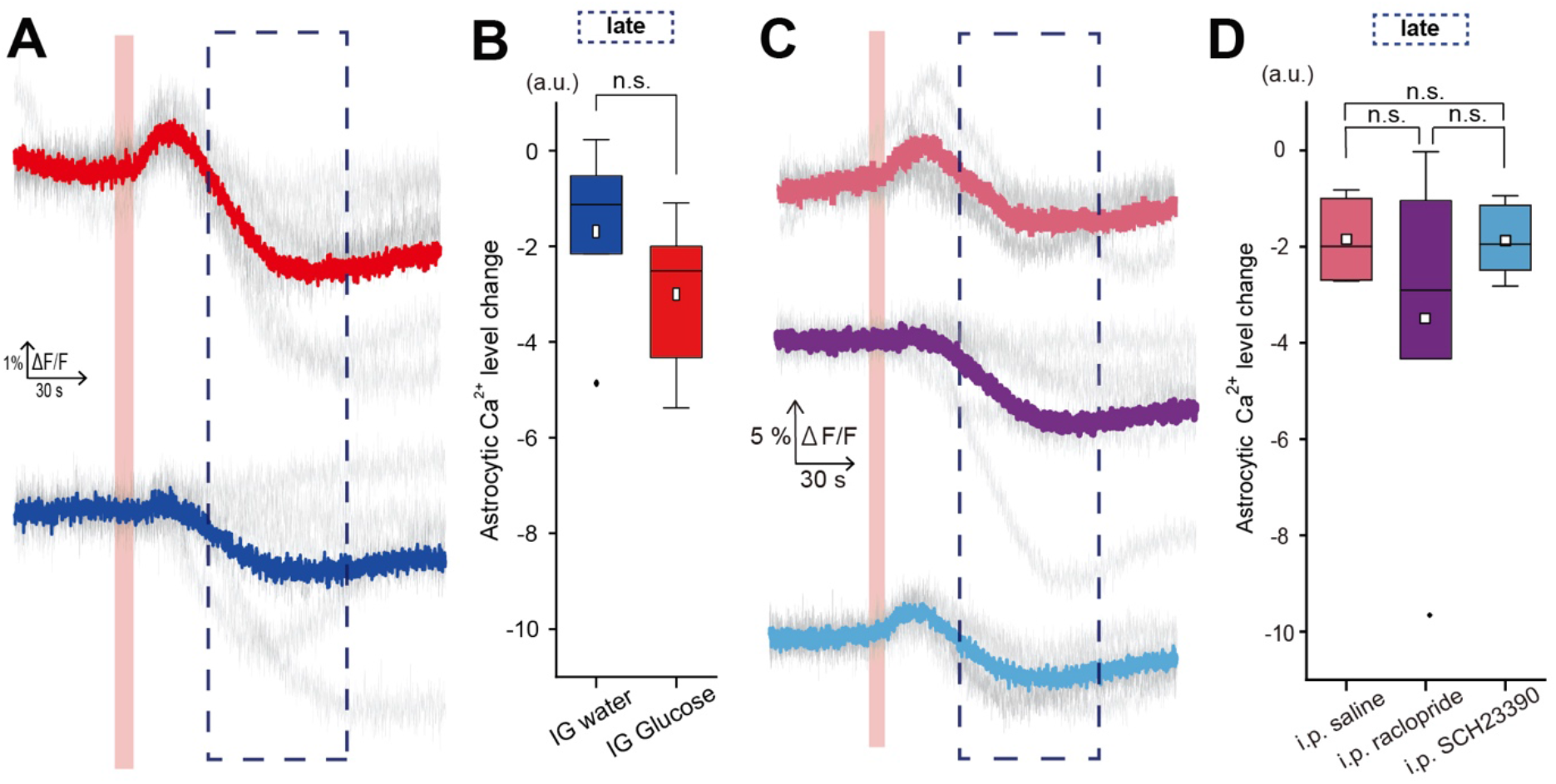
Late-phase astrocytic signal dynamics in M2 under IG glucose and water injections. (A) Traces of averaged Mlc-YC (Astrocytes) signal responses to IG glucose injection (red bold line) and IG water injection (blue bold line), with the light gray line depicting traces from individual subjects. (glucose; n = 7 mice, water; n = 5 mice) (B) Comparative analysis of late-phase changes in Mlc-YC (Astrocytes) signals in response to IG water injection (blue) and IG glucose injection (red). The analysis window (late phase) corresponds to the area within the dotted line in (A). (water; N = 5 mice, glucose; N = 5 mice) (C) Traces of averaged Mlc-YC (Astrocytes) signal responses to IG glucose injection following i.p. saline (pink bold line), raclopride (purple bold line), and SCH23390 (light blue bold line) injections, with the light gray line representing traces from individual subjects. (saline; N = 5 mice, raclopride; N = 8 mice, SCH23390; N = 7 mice) (D) Comparative analysis of late-phase changes in Mlc-YC (Astrocytes) signals in response to IG glucose injection following i.p. saline (pink), raclopride (purple), and SCH23390 (light blue) injections. The analysis window (late phase) corresponds to the area within the dotted line in (C). (saline; N = 5 mice, raclopride; N = 8 mice, SCH23390; N = 7 mice)

**Supplementary Fig. 3.**
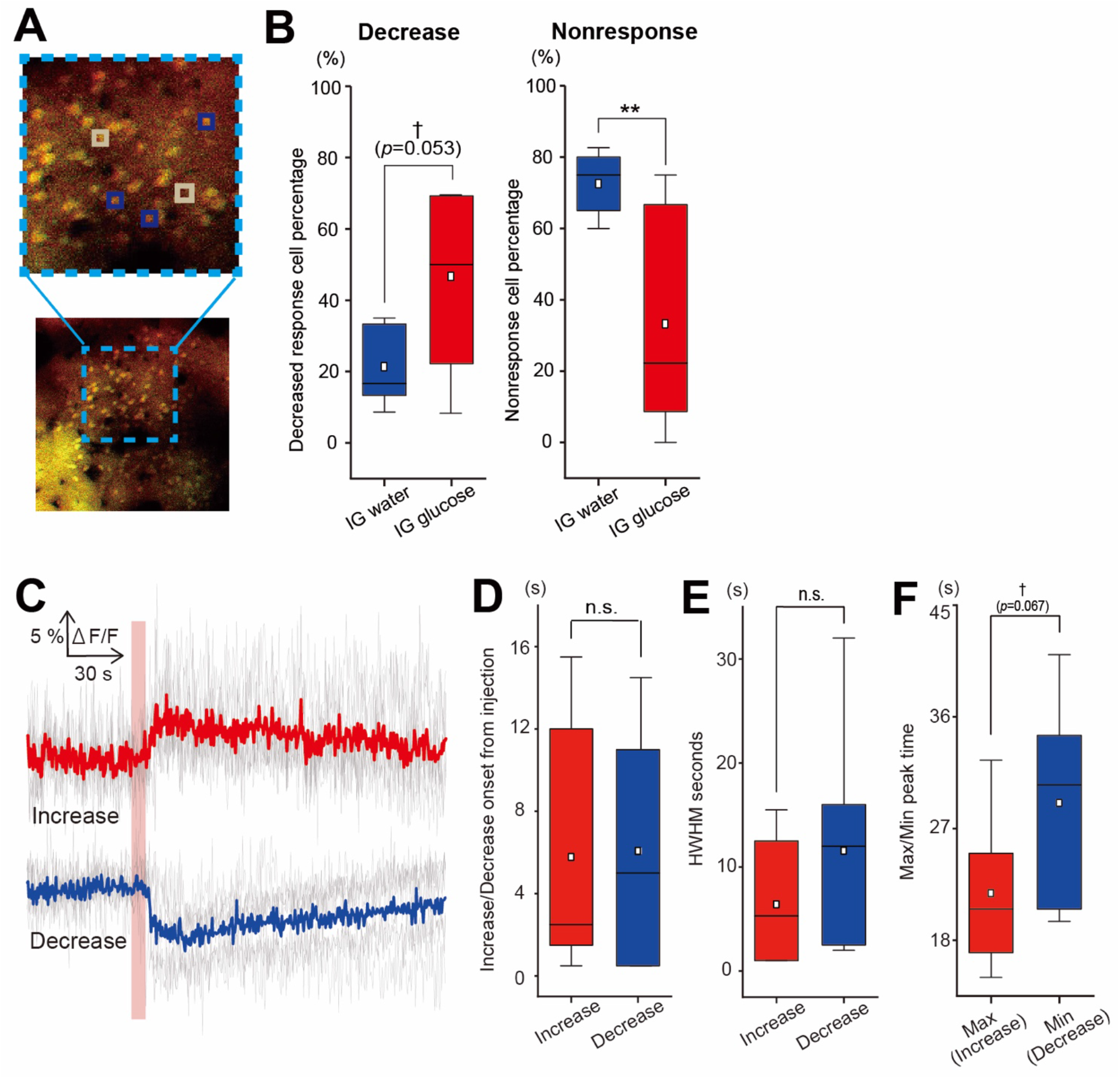
Late-phase single cell level astrocytic signal dynamics in M2 under IG glucose injections. (A) Representative two-photon microscopy image of layer 2 in the frontal cortex, with cells labeled with Fluo4-AM (green; Ca^2+^ indicator) and SR101 (red; astrocytes). Colored squares mark regions of interest (ROIs); cells encircled by blue squares exhibited a decrease in reaction, while those surrounded by beige squares showed no response. (B) Comparative analysis of the proportion of cells displaying a decrease and no response among all evaluated cells for each subject. **p<0.01, †<0.1 (IG glucose; N = 7 mice, IG water; N = 5 mice, two-sample t-test) (C) ΔF/F traces of fluorescence intensity for the cell populations identified to exhibit an increase and decrease under IG glucose injection (gray line) alongside the corresponding aggregate data (bold red and blue lines, respectively). (Increase; 25 cells from N = 7 mice, Decrease; 69 cells from N = 7 mice) (D) Comparative analysis of the onset times for increases and decreases under IG glucose injection. (N=7 mice, two-sample t-test) (E) Comparative analysis of the Half-Width at Half Maximum (HWHM) for the average data of each subject identified to exhibit an increase and decrease. (N = 7 mice, two-sample t-test) (F) Comparison of the Max and Min peak time of the fluorescence intensity of cells which exhibit increase and decrease respectively, under IG glucose injection. †<0.1 (N = 7 mice, two-sample t-test)

